# Bionized nanoferrite particles alter the course of experimental *Cryptococcus neoformans* pneumonia

**DOI:** 10.1101/2021.12.21.473767

**Authors:** Livia C. Liporagi Lopes, Preethi Korangath, Samuel R. Dos Santos Junior, Kathleen L. Gabrielson, Robert Ivkov, Arturo Casadevall

## Abstract

Cryptococcosis is a devastating fungal disease associated with high morbidity and mortality even when treated with antifungal drugs. Bionized nanoferrite (BNF) nanoparticles are powerful immunomodulators, but their efficacy for infectious diseases has not been investigated. Administration of BNF nanoparticles to mice with experimental cryptococcal pneumonia altered the outcome of infection in a dose response manner as measured by colony forming units and survival. The protective effects were higher at lower doses, with reductions in IL-2, IL-4 and TNF-α, consistent with immune modulation whereby reductions in inflammation translate into reduced host damage, clearance of infection and longer survival.

## Introduction

*Cryptococcus neoformans* is a fungal pathogen with a worldwide distribution. Serological studies of human populations show a high prevalence of human infection, which rarely progresses to disease in immunocompetent hosts (1, 2). However, decreased host immunity places individuals at high risk for cryptococcal disease. The disease can result from acute infection or reactivation of latent infection, in which yeasts within granulomas and host macrophages emerge to cause disease (3). In both immunocompetent and immunosuppressed patients, cryptococcosis has high morbidity and mortality, even with aggressive antifungal drug therapy (1). Current therapies can require anti-fungal drugs for several months and treatment can be complicated by rising antimicrobial drug resistance (4, 5). Consequently, development of new therapeutics against *C. neoformans* infections is urgently required.

Magnetic iron oxide nanoparticles (MNPs) have proven useful for the diagnosis and therapy of various conditions because they display generally favorable biocompatibility and varied responsiveness to magnetic fields (6–10). When injected into the bloodstream, nanoparticles encounter a complex fluid environment that can modify the initial particle surface to one having a molecular signature which produces specific interactions with host biology (11–13). Ultimately, though indirectly, the interactions of nanoparticles with the complex host environment can lead to their deposition in organs or tissues that depends upon the initial physical and chemical properties of the injected construct (14). The immune system’s function in the maintenance of tissue homeostasis is to protect the host from environmental agents such as microbes or chemicals, and thereby preserve the integrity of the body. The study of the interactions between nanoparticles and various components of the immune system is an active area of research in bio- and nanotechnology because the benefits of using nanotechnology in industry and medicine are often questioned over concerns regarding the safety of these novel materials (11, 15, 16). The past decade of research has shown that, while in certain situations nanoparticles can be toxic, nanotechnology engineering can modify these materials to either avoid or target the immune system. Depending on the nanoparticle composition and physicochemical properties, nanoparticle interactions with host biology can induce or increase inflammation (17, 18) or mediate immunosuppression (19, 20) Either of these responses can be used to enhance efficacy in specific disease contexts, and which response is desired depends on specific features of both the disease and nanoparticle. For a nanoparticle to induce both inflammation and immunosuppression in the same disease context however is unusual. Specific targeting of the immune system, on the other hand, provides an attractive option for vaccine delivery, as well as for improving the quality of anti-inflammatory, anticancer, and antiviral therapies (21–24). Macrophages utilize multiple routes to ingest the same types of nanoparticles (25). Several studies reported that smaller particles (20–200 nm) elicit stronger immune responses than their larger counterparts (11, 26–29).

Previous studies have shown that injection of nanoparticles into experimental animals elicited pseudo infection-like response or local inflammation or systemic immune response leading to immunological changes in tumor microenvironment in mouse models of caner (30, 31). When delivered systemically, BNF nanoparticle can trigger an immune response that resembles infection, leading to downstream signaling that engages effector anti-tumor CD8^+^ T cell function (31). The effect of nanoparticles on tumor biology has attracted considerable attention but there has been no comparable effort to study their effect on infectious diseases,. Like other engineered nanoparticles (NPs), BNF NPs inherently possess physical features resembling viruses and elicit an antitumor immune response when injected into immune competent mouse models of breast cancer (31).

In this study we tested the effect of BNF in a mouse model of cryptococcal pneumonia. Our results show that BNF nanoparticles have a profound effect on the inflammatory response to *C. neoformans* infection that is associated with altered outcomes depending on the nanoparticle dose used.

## Results

The fungal burden was assessed from the removed lungs and mice treated with BNF at high dose had higher CFUs (**Figure 1A)**; CFU numbers between groups were not statistically significant, but there was a trend towards higher fungal burden in the BNF-treated groups, with higher numbers in groups that received BNF 24 h after the challenge with *C. neoformans* (**Figure 1A**). Histological slides of lungs showed that PBS treated group had some inflammation (**Figure 1B**) whereas nanoparticle injection 5mgFe/mouse) before (**Figure 1C**) or after 24 h (**Figure 1D**) of infection had increased inflammation seen by H&E. It is possible to observe in the slides an increased number of yeasts in both nanoparticle-treated groups (**Figures 1F and 1G**) compared to control (**Figure 1E**) as shown by PAS staining of lungs. This aggravated reaction could be due to the very high concentration of nanoparticle used in this initial study.

**Figure 1:**
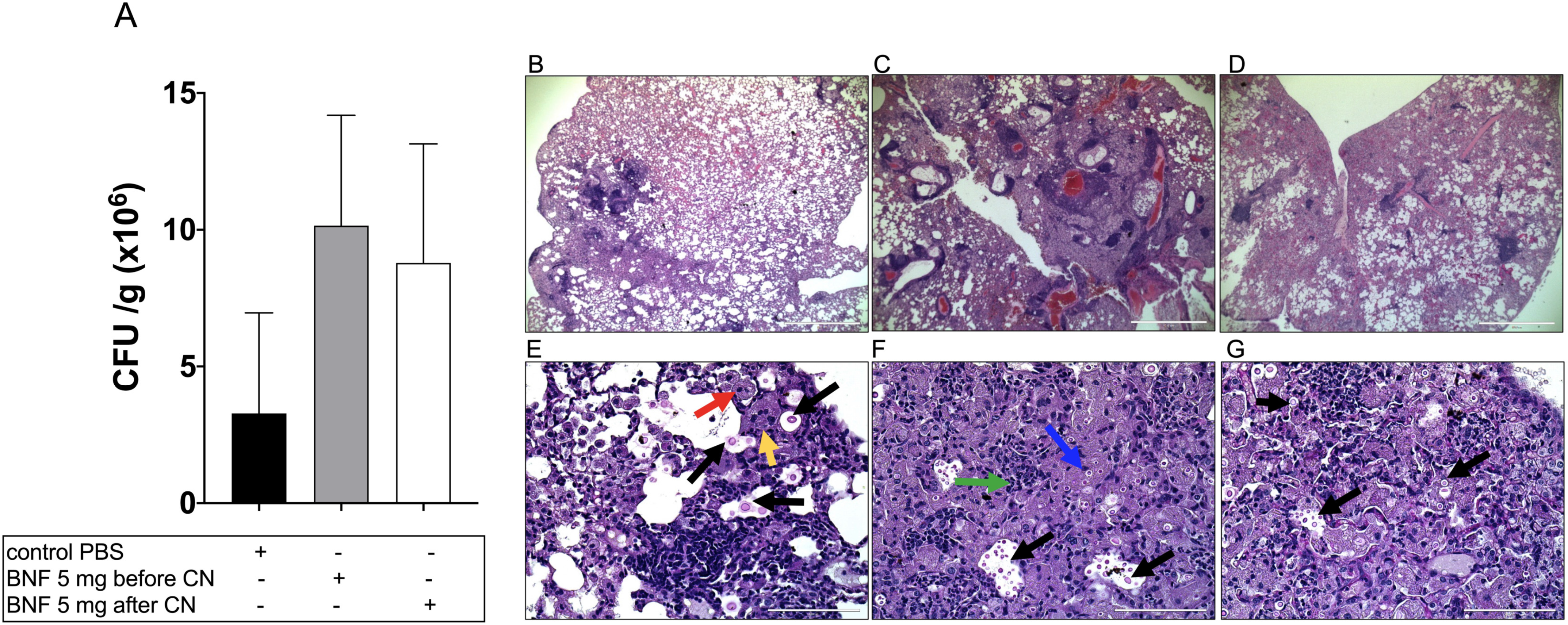
Higher recovery of lungs CFU 60 days after challenge with *C. neoformans* strain 24067 from the BNF-treated groups compared with the PBS control group (**A**). Intense immune reaction with numerous yeast organisms was seen following treatment with higher concentrations of nanoparticles in FVB/NJ mice. Representative histology images after 60 days of infection in PBS treated group (**B**), nanoparticle injection (5mgFe/mouse) before 24hrs (**C**) or after 24hrs (**D**) by H&E. It is possible to observe an increased number of yeast in both nanoparticle treated groups (**F, G**) compared to control (**E**) by PAS. Black arrows in **E-G** indicate yeast, red arrows indicates foamy macrophages and yellow indicates multinucleated giant cells. Blue arrow in **F** indicates organism inside a macrophage and green indicates neutrophils.

Given the results from the high dose BNF experiment in FVB/N mice, we evaluated whether lower doses of BNF nanoparticles in the same model of FVB/N mice would produce different outcomes. A significant decrease (p<0.05) of fungal CFU from the recovered lungs was observed in mice treated with lowest dose of BNF (BNF 0.005 mg) (**Figure 2A**), demonstrating that at lower doses, BNF NPs (as 0.005 mg) could protect against *C. neoformans* infection. Corroborating with CFU data, H&E stained lung sections show inflammatory cells surrounding Cryptococcus yeast in FVB/NJ mice treated with PBS, with numerous macrophages and multinucleated giant cells (blue arrow) and neutrophils (green arrow **Figures 2B and 2C**). BNF (0.005 mg) treated lungs had fewer inflammatory cells (**Figures** 2**D** **and 2E**). PAS staining showed presence of multiple fungal yeasts in PBS treated mice (**Figure 2F**) whereas lower concentration (BNF 0.005 mg) of nanoparticle treated lungs showed no positive staining for yeast (**Figure 2G**). Prussian blue staining for nanoparticles in lungs also showed no evidence of iron (BNF) deposits (data not shown).

**Figure 2:**
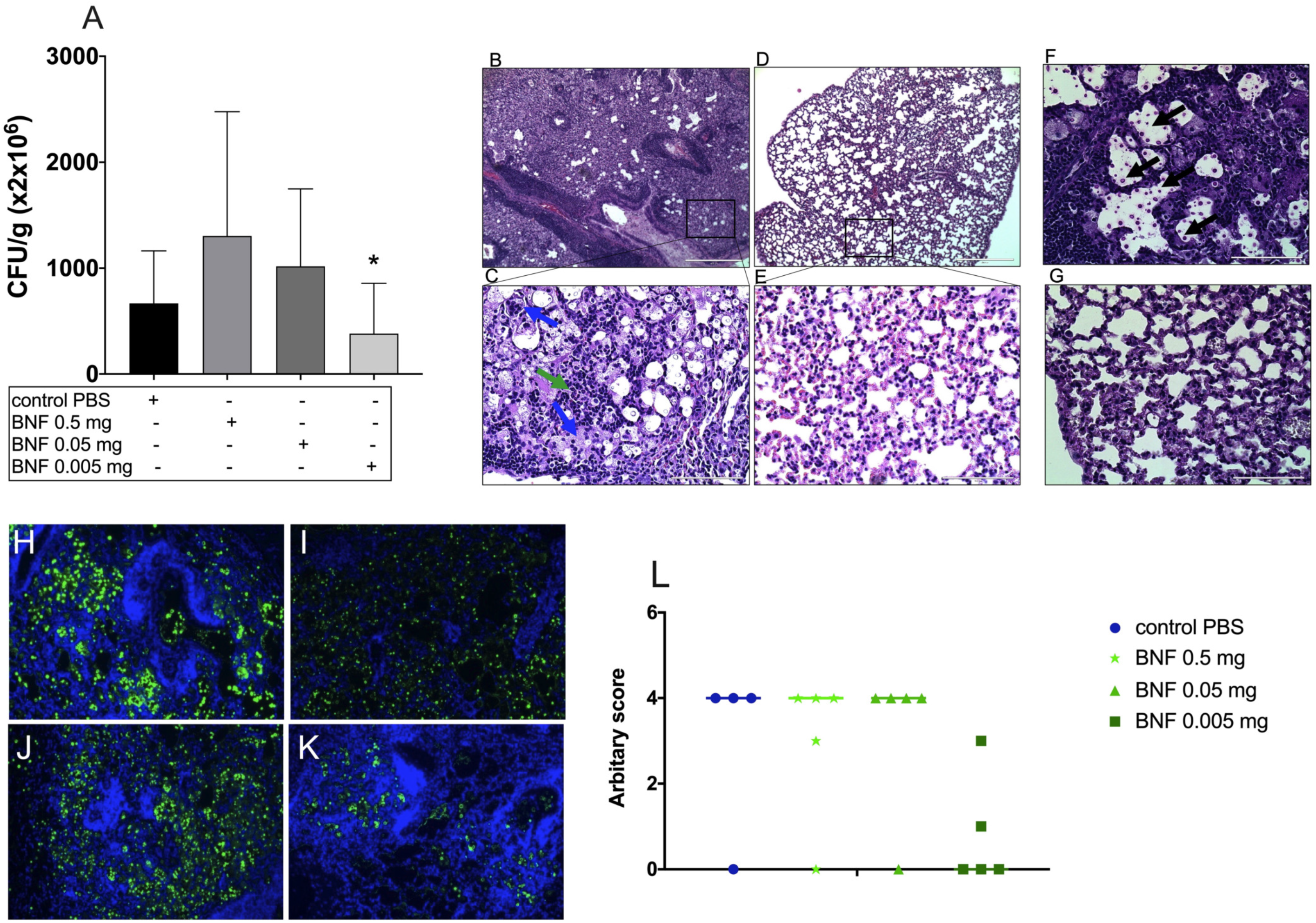
BNF 0.005 mg doses showed a significant decrease (p<0.05) of fungal CFU from recovered lungs (**A**) indicating a protective profile in lungs 14 days after challenge with *C. neoformans* strain 24067. H&E stained lung in FVB/NJ mice treated with PBS showing numerous macrophages and multinucleated giant cells (blue arrow) and neutrophils (green arrow) (**B, C**). BNF (0.005mg) treated lungs (**D, E**). PAS staining in PBS treated mouse (**F**) and in nanoparticle treated lungs (**G**). Tissues were stained with monoclonal antibody 18B7 conjugated with Oregon green with DAPI. Control PBS (**H**), BNF 0.5 mg (**I**), BNF 0.05 mg (**J**), BNF 0.005 mg (**K**). 10X magnification. Representative graph showing the IF-score generated by the IF slides (**L**).

To further analyse the protective effect observed in FVB/N mice treated with lower dose of BNF, we carried out a survival study in A/J mice. A/J mice were chosen to test whether the high innate resistance of FVB/N mice to cryptococcal infection, which survived a high *C. neoformans* inoculum such that none died after 60 days, was responsible for the observed effects. In this experiment, we analysed both mouse survival and lung fungal burden in different groups. Mice receiving the lowest dose of (0.0005 mg) showed a significant increase (p<0.05) in the survival, with no deaths (survival of 100%) after 60 d, in comparison to the control group (PBS injected) with a survival of only 20%. (**Figure 3A**). To analyse the fungal burden, lungs were removed 14 days after infection and CFU numbers were determined. Similarly, a significant decrease (p<0.05) in the number of recovered lung CFU was observed in the group that received the BNF 0.0005 mg (**Figure 3B**). These data suggest that at lower dose, systemic treatment with BNF NPs can stimulate a protective response when challenged by two different strains of *C. neoformans* in two different mouse strains. We observed nearly complete remission of disease following a single treatment with low dose BNF NPs in A/J mice. Analysis of H&E stained lung sections revealed numerous and varied inflammatory cells surrounding *Cryptococcus* yeast indicative of a robust and organized immune response (blue arrow) (**Figures 3C and 3D**). Additionally, BNF nanoparticle (0.0005 mg Fe/mouse) treated mouse lungs had fewer inflammatory cells in lungs (**Figures 3E and 3F**). PAS staining showed presence of numerous *Cryptococcus* in PBS treated control mouse lung (**Figure 3G, black arrow**). BNF nanoparticle (0.0005 mgFe) treated mouse lungs showed no positive staining for yeast (**H**). We noted less mAb 18B7 immunofluorescence in mouse tissues treated with low dose nanoparticles (0.0005 mg) (**Figures 3K and 3L**). Mice treated with higher nanoparticle doses (0.005 mg) (**Figures 3J and 3L**) had *C. neoformans* similar to that of control group (**Figures 3I and 3L**). In the lungs and spleens of mice given the lower doses of BNF the levels of the proinflammatory cytokines tumor necrosis factor-alpha (TNF-α) was decreased after 14 days of infection on BNF 0.005 mg group relative to control mice infected with *C. neoformans* (p<0.05)(**Figure 3M**). Additionally, interleukin-2 (IL-2), interleukin-4 (IL-4), interleukin-10 (IL-10) and interferon-gamma (IFN-γ) were analysed but no statistically differences were observed between the groups (**Figure 3M**). We focused on the low BNF doses since these were associated with better control of infection.

**Figure 3:**
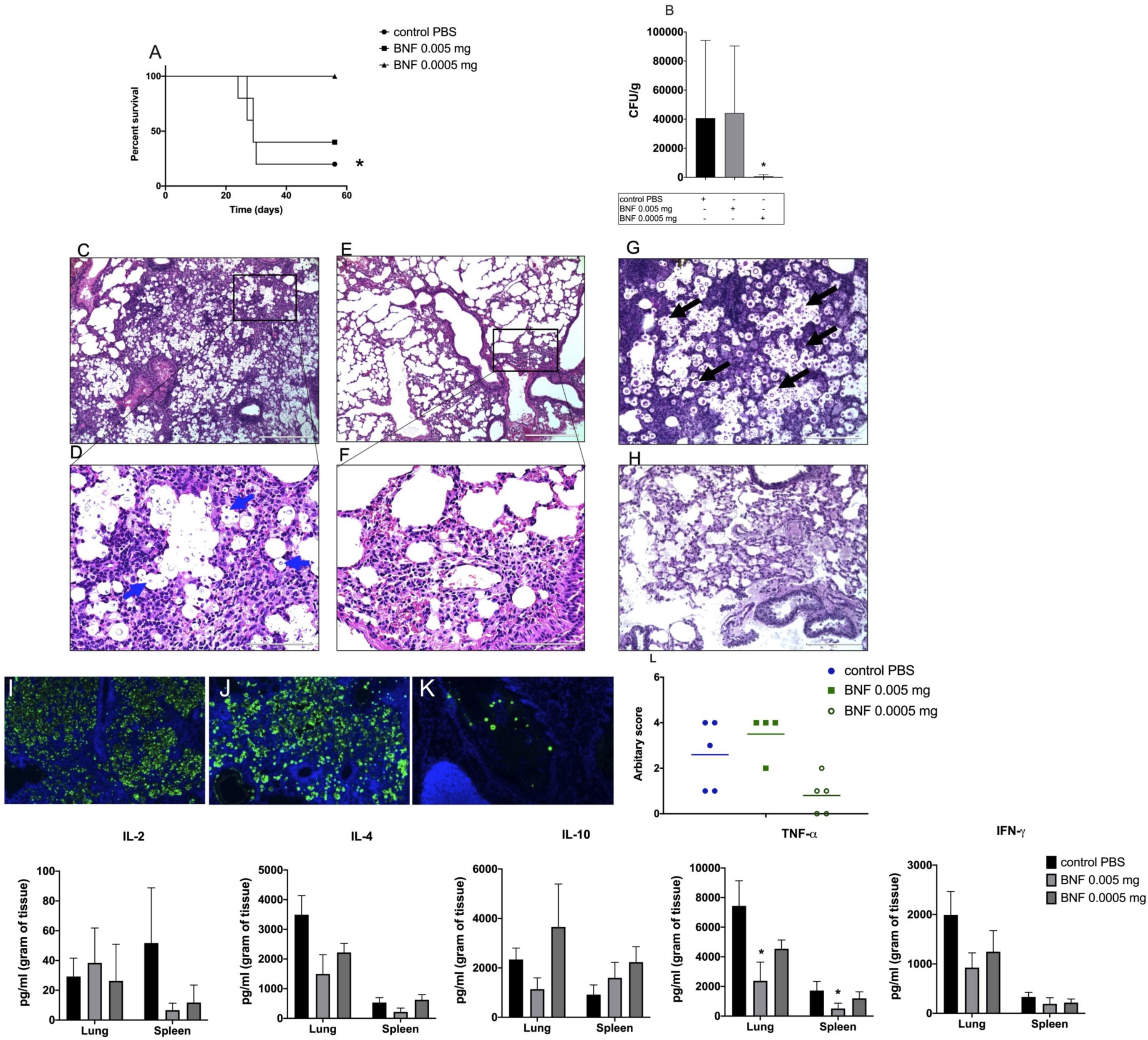
A/J mice injected with lower doses of BNF showed a protective profile in lungs 14 days after challenge with *C. neoformans* strain H99. Mice injected with the lower doses of BNF nanoparticles (0.0005 mg) showed a survival of 100% in comparison to the control group (no BNF nanoparticle injection) with a survival of only 20% (p<0.05) (**A**). Lung fungal burden showed a significant decrease (p<0.05) in the number of recovered CFU in the mice that received the BNF 0.0005 mg (**B**). H&E stained lung sections showed numerous inflammatory cells surrounding *Cryptococcus* yeast (blue arrow) (**C, D**). BNF nanoparticle (0.0005mgFe/mouse) treated mouse (**E, F**). PAS staining showed presence of numerous *Cryptococcus* in PBS treated control mouse lung (**G**, black arrow). BNF nanoparticle (0.0005mgFe) treated (**H**). Tissues were stained with monoclonal antibody 18B7 conjugated with Oregon green with DAPI (10x magnification). Control PBS (**I**), BNF 0.005 mg (**J**), BNF 0.0005 mg (**K**). Representative graph showing the IF-score generated by the IF slides (**L**). Lung and spleen cytokine levels during the course of infection (**M**). *p<0.05.

Lab culture passages can affect the pathogenicity of cultured *Cryptococcus* strains (32). To test the potential effects of lab strain variability, we thawed a second vial of strain H99 to infect mice in Study IV to compare the effects with assays of fungal burden and cytokines from lung and spleen in A/J mice. In contrast to results obtained from Study III (**Figure 3B**), there was no statistically significant difference in the number of recovered CFUs measured in lungs between PBS control and BNF 0.0005 mg groups (**Figure 4A**). There was, however, a slight decrease in the CFUs number measured in the BNF 0.0005 mg Fe group from spleen, but this was not statistically significant (**Figure 4A**). From the lung and spleen cytokines analysis, levels of proinflammatory cytokines, including interleukin-2 (IL-2), interleukin-4 (IL-4), tumour necrosis factor-alpha (TNF-α) and interferon-gamma (IFN-γ) were decreased after 14 days of infection relative to control mice infected with *C. neoformans* (**Figures 4B**), but this difference was statistically significant only for IL-2 (p<0.005). Interleukin-10 (IL-10) and interferon-gamma (IFN-γ) were analysed but no statistically differences were observed between both groups (**Figure 4B**).

**Figure 4:**
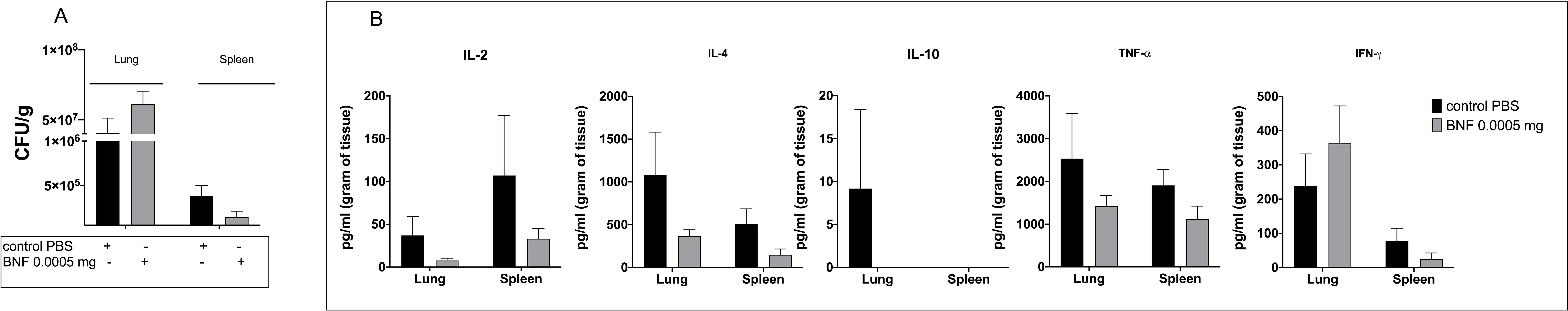
Fungal burden from lungs and spleens of A/J mice 14 days after challenge with recently thawed *C. neoformans* strain H99 showed a slightly decrease from spleens of BNF 0.0005 mg group compared with PBS control groups (**A**). Lung and spleen cytokines levels were also analysed (**B**).

**Table 1** summarizes all the parameters accessed and analysed between Study I - Study 4. The four independent experiments reported here each differs in several variables and they represent our experience as we modified the conditions sequentially in an effort to find a set of parameters that resulted in benefit to the mice. Clearly, changes in such variables as mouse background, inoculum, cryptococcal strain and BNF dose can each have large effects in the outcome of individual experiments. Hence, we caution about interpretations from individual experiments and instead stress that the common result across all the experiments was that BNF administration affected the outcome of *C*. *neoformans* infection in mice. These exploratory studies provide encouragement for future additional work to delineate the conditions where BNF particles can be used to modify the outcome of infection.

**Table 1:** Summary of all the parameters from Study 1, Study 2, Study 3 and Study IV.

| Experiment | BNF doses | Mice strain | Number of mice | Fungal strain | Time of mice infection | Parameters assessed | CFU data (LUNG) | CFU data (SPLEEN) | Survival curve (60 days) | Histology (LUNG) | IF (LUNG) | IL-2 | IL-4 | IL-10 | IFN-γ | TNF-α |
| --- | --- | --- | --- | --- | --- | --- | --- | --- | --- | --- | --- | --- | --- | --- | --- | --- |
| Study I | 5 mg (24 h after and before fungal challenge) | FVB/N | 5 per group | 24067 | 60 days | CFU, histology (H&E, PAS) | no difference between groups | X | no difference between groups | no difference between groups | X | X | X | X | X | X |
| Study II | 0.5/0.05/0.005 mg | FVB/N | 5 per group | 24067 | 14 days | CFU, histology (H&E, PAS, IF) | lower CFU number on 0.005 mg (p<0.05) | X | X | less yeasts and inflammation on 0.0005 mg mice group | less yeasts on 0.0005 mg mice group | X | X | X | X | X |
| Study III | 0.005/0.0005 mg | A/J | 10 per group | H99 | 14 days to sacrifice / 60 days survival | CFU, histology (H&E, PAS, IF), survival, cytokine | lower CFU number on 0.0005 mg (p<0.05) | X | higher survival on both 0.005 & 0.0005 mg groups, (P<0.05 on 0.0005 mg group) | less yeasts and inflammation on 0.0005 mg mice group | less yeasts on 0.0005 mg mice group | tendency to decrease (lung) | tendency to decrease (lung) | no difference | tendency to decrease (lung and spleen) | decreased (lung and spleen) on 0.005 mg group (p<0.005) |
| Study IV | 0.0005 mg | A/J | 10 per group | H99 | 14 days | CFU, cytokine | no difference between groups | lower CFU number on 0.0005 mg | X | X | X | decreased (lung and spleen) on 0.0005 mg group (p<0.005) | tendency to decrease (lung and spleen) | no difference | tendency to decrease (lung and spleen) | tendency to decrease (lung and spleen) |

## Discussion

Clinically, Cryptococcosis is most prevalent in immunocompromised patients but a significant proportion of cases occur in patients with no apparent immune deficit. One of the confounding paradoxes of this disease is that the prognosis is better in patients with AIDS, possibly because host damage is a function of both fungal processes such as increased intracranial pressure and also tissue damaging effects inflammation (reviewed in (33)). Consequently, a common denominator in human cryptococcosis, whether it occurs in immunosuppressed or immunologically intact individuals is dysregulated inflammatory responses, that are ineffective to contain the infection. Since deficits in immune function appear to be major contributors to the outcome of cryptococcosis it is reasonable to posit that future improvements in therapy will require immune modulators, rather than immune stimulating or immune suppressing treatments (34). In the present study BNF nanoparticles altered the outcome of infection in a dose response manner, with the best effects observed with low nanoparticle doses, consistent with down-regulation of the inflammatory response, possibly associate with immune modulation. Higher doses of BNF particles were associated with stronger inflammation and a trend toward higher fungal burden, suggesting they triggered exuberant immune responses that aggravated infection and damaged host tissue. The association of stronger inflammatory response with lower efficacy is consistent with emerging evidence that host damage in cryptococcosis can have a strong immune mediated component (33).

In summary, our results show that administration of BNF nanoparticles before and after experimental cryptococcal pneumonia infection in mice alters the outcome of infection. Taken together, these preliminary results indicate that iron oxide nanoparticles may harbour immune modulating capabilities to increase/induce inflammation and to suppress/reduce, depending on dose. In this context, nanoparticles comprise important characteristics that make them attractive for a variety of biomedical applications (35). Specifically, iron oxide nanoparticles are physically and chemically stable, biocompatible, and environmentally safe (35), thus presenting unique characteristics for clinical applications. To our knowledge this is the first exploration of BNF nanoparticles in an infectious disease model and our results establish the proof of principle that BNF nanoparticles can affect the course of infectious diseases in a manner similar to the experience with cancer and inflammatory diseases. Our results encourage further studies to determine whether they can be used for immunotherapy of cryptococcosis as an adjuvant of standard antifungal therapy.

## Materials and Methods

*C. neoformans* serotype D strain 24067 was used in all FVB/N mouse experiments. *C. neoformans* var *grubii* serotype A strain H99 (ATCC 208821) was used in all A/J mice. The yeast cells were kept frozen in 10%-glycerol. Sabouraud dextrose broth (SAB, from Gibco) medium was used for standard growth of yeast cells at 30 °C with moderate shaking (120 rpm) overnight.

BNF NPs are commercially available as aqueous suspensions of hydroxyethyl starch-coated magnetite (Fe_3_O_4_) core-shell nanoparticles (Micromod Partikeltechnologie, GmbH, Rostock, Germany). Their synthesis and physical and biological characterization has been extensively documented by us (31, 36–39). BNF NPs are synthesized by precipitating ferric and ferrous sulfate salts from solution at high pH in a high-pressure-homogenization reaction vessel. According to the manufacturer, they have a mean hydrodynamic diameter of ~100 nm (polydispersity index < 0.25); approximately neutral zetapotential (~-2 mV @ pH=7.4); and, Fe content >50% w/w (or iron oxide >70% w/w) (31).

All animal procedures were performed with prior approval from Johns Hopkins University (JHU) Animal Care and Use Committee (IACUC), under approved protocol number MO18H152. JHU Animal Welfare Assurance Number is D16-00173 (A3272-01). Two mouse strains were used to perform the experiments: six-week-old female A/J mice (from Jackson Laboratory - JAX stock #000646) and six- to eight-week-old female FVB/N (from Jackson Laboratory - JAX stock #001800). A/J mice were used because they are highly susceptible to cryptococcal infection (40) whereas our use of FVB/N mice was motivated by our previously reported findings of anti-cancer immune modulation by BNF NPs in this mouse strain [31]. Four different murine experiments were performed as described below. In all experiments, each mouse received a single intravenous (i.v.) injection of either PBS (150 μl/mouse) or BNF at the described dose, according to its group assignment:

Experiment I –Fifteen female FVB/N mice were divided into three groups (n = 5 animals per group): Group 1 –PBS 24 h after *C. neoformans* infection (control PBS); Group 2 – BNF 5 mg Fe in 150 μl/mouse 24 h after *C. neoformans* infection (BNF 5 mg after CN); and Group 3 – BNF 5 mg Fe in 150 μl/ mouse) 24 h before *C. neoformans* infection (BNF 5 mg before CN). Sixty days after infection, surviving animals were euthanized and tissues extracted for fungal burden and histology analysis.

Experiment II – Experimental design was modelled from Study I, but with de-escalating BNF dose to determine dose-response. Analysis of Study I results indicated potent immune responses, motivating a dose de-escalation study. Another twenty female FVB/N mice were divided into four groups (n = 5 animals per group), representing PBS control, and BNF dose: Group 1 – PBS 24 h before *C. neoformans* infection (control PBS); Group 2 – BNF 0.5 mg Fe in 100 μl/animal 24 h before *C. neoformans* infection (BNF 0.5 mg); Group 3 – BNF 0.05 mg Fe in 100 μl/animal 24 h before *C. neoformans* infection (BNF 0.05 mg); and Group 4 – BNF 0.005 mg Fe in 100 μl/animal 24 h before *C. neoformans* infection (BNF 0.005 mg). Animals were observed daily for 14 days and surviving animals were euthanized, and tissues extracted for fungal burden and histology analysis on 14 days after of infection.

Experiment III – Study II results showed lower dose of BNF nanoparticles protected FVB/N mice from *C. neoformans* infection, motivating further study in the A/J strain with established susceptibility to *C. neoformans* infection. Thirty A/J mice (n = 10 animal per group) were divided into three groups: Group 1 – PBS 24 h before *C. neoformans* infection (control PBS); Group 2 – BNF 0.005 mg Fe in 100 μl/animal 24 h before *C. neoformans* infection (BNF 0.005 mg); and Group 3 – BNF 0.0005 mg Fe in 100 μl/animal 24 h before *C. neoformans* infection (BNF 0.0005 mg). Five animals from each group were euthanized 14 days after infection for fungal burden, histology and cytokine analysis and the other five were followed for survival until 60 days.

Experiment IV - Another cohort of twenty A/J mice (n = 10 animal per group) were divided into two groups: Group 1 – PBS 24 h before *C. neoformans* infection (control PBS); and Group 2 – BNF 0.0005 mg Fe in 100 μl/animal 24 h before *C. neoformans* infection (BNF 0.0005 mg). All animals were sacrificed on day 14 for fungal burden and cytokine analysis from lung and spleen.

All mice challenged with *C. neoformans* were intranasally infected with 1 x 10^7^ *C. neoformans* yeasts per animal, in a total volume of 20 μl (10 μl in each nasal cavity of the mouse). This model produces a pulmonary infection that rapidly disseminates to brain and other organs (41). Mice were anesthetized with 60 μl xylazine/ketamine solution intraperitoneally (95 mg of ketamine and 5 mg of xylazine per kilogram of animal body weight) to perform intranasal infection.

The fungal burden in lungs was measured in surviving mice by counting colony-forming units (CFU) (41, 42). At the endpoint of each experiment (as described above), mice were euthanized and the lungs (left lobe) were removed. Organ sections were weighed and homogenized in 1 ml of PBS. After serial dilutions, homogenates were inoculated on Sabouraud agar plates with 10 U/ml of streptomycin/penicillin. The plates were incubated at room temperature, and colonies counted after 48-72 h.

A piece of the right lung and a piece of the spleen were fixed in 10% formalin for 48 h before embedding in paraffin. Tissue slides were stained with Hematoxylin and Eosin (H&E), Periodic acid-Schiff (PAS) for *C. neoformans* and Prussian blue to visualize nanoparticles. The remaining right lungs and spleens of each A/J mouse treated with PBS or BNF 0.0005mg were macerated with protease inhibitor (complete, EDTA-free, Roche Life Science, IN, USA) and centrifuged; supernatants of these samples were used for cytokine detection by sandwich-ELISA with commercial kits (BD OptEIA™, BD Franklin Lakes, NJ, USA) for the following cytokines: IL-2 (#555148), IL-4 (#555232), IL-10 (#555252), IFN-γ (#551866) and TNF-α (#555268).The protocol was followed according to the manufacturer’s recommendations. Readings were performed in a plate spectrophotometer at 450 and 570 nm.

Immunofluorescence (IF) staining was performed on lung tissues to analyse the capsular polysaccharide covering *Cryptococcus* yeasts. Briefly, slides were deparaffinised on a slide warmer at 58°C 10 minutes followed by serial washing in 2 changes of xylene, 100%, 95% and 70% alcohol. Slides were dehydrated in deionized water and then treated with the antibody in a blocking solution for 30 minutes. The slides were then stained with the antibody (18B7) (43) conjugated with Oregon green during 16h at 4°C. On the next day all slides were washed with at least 5 changes of PBS and mounted with DAPI containing mounting media (ProLong Gold Antifade Mountant - (ThermoFisher Scientific, MA, USA). Slides were then visualized and imaged with a Zeiss microscope with 100X magnification.

*S*tatistical analyses were done using GraphPad Prism version 8.00 for Mac OS X (GraphPad Software, San Diego, CA, USA). One-way analysis of variance using a Kruskal-Wallis nonparametric test was used to compare the differences between groups, and individual comparisons of groups were performed using a Bonferroni post-test. The Student’s t-test was used to compare the number of colony forming units (CFU) for different groups. The 90–95% confidence interval was determined in all experiments. Unpaired t-test (F test to compare variances) was performed to compare cytokines data.

## Acknowledgments

AC was supported in part by NIH grants AI052733, AI15207 and HL059842. This work was supported in part by Johns Hopkins Discovery Award.

